# BIGMAC: Breaking Inaccurate Genomes and Merging Assembled Contigs for long read metagenomic assembly

**DOI:** 10.1101/045690

**Authors:** Ka-Kit Lam, Richard Hall, Alicia Clum, Satish Rao

## Abstract

The problem of de-novo assembly for metagenomes using only long reads is gaining attention. We study whether post-processing metagenomic assemblies with the original input long reads can result in quality improvement. Previous approaches have focused on pre-processing reads and optimizing assemblers. BIGMAC takes an alternative perspective to focus on the post-processing step. Using both the assembled contigs and original long reads as input, BIGMAC first breaks the contigs at potentially mis-assembled locations and subsequently scaffolds contigs. Our experiments on metagenomes assembled from long reads show that BIGMAC can improve assembly quality by reducing the number of mis-assemblies while maintaining/increasing N50 and N75. The software is available at https://github.com/kakitone/BIGMAC

## 1 Introduction

De-novo assembly is a fundamental yet difficult [1] computational problem in metagenomics. Indeed, there is currently an open challenge for metagenomic assembly using short reads, titled “Critical Assessment of Metagenomic Interpretation (CAMI [2]).” On the other hand, emerging sequencing technologies can provide extra information and make the computational problem more tractable. For example, long reads are increasingly being shown to be valuable in the de-novo assembly of single genomes[3]. Therefore, it is natural to study metagenomic assembly using long reads. Current assembly approaches for long reads focus on developing more optimized assemblers to leverage the power of the data. However, significant engineering effort is usually involved to build an end-to-end assembler.

We take a different perspective, focusing the design effort on a post-processor that improves assembled contigs using the original long read data (Fig 1). The main question is whether we can achieve quality improvement with this approach using typical long reads. This post-pocessing approach is attractive because it leverages existing software. Consequently, we can focus design effort and computational resources to specifically address thorny issues arising from the nature of new data, complex repeats, varying abundances and noise. Moreover, since the long reads are part of the input, the post-processor can bring back information missed upstream. In single genome assembly, FinisherSC [4] has demonstrated the effectiveness of this approach. In this paper, we investigate the effectiveness of this post-processing approach for metagenomic assembly.

**Figure 1:**
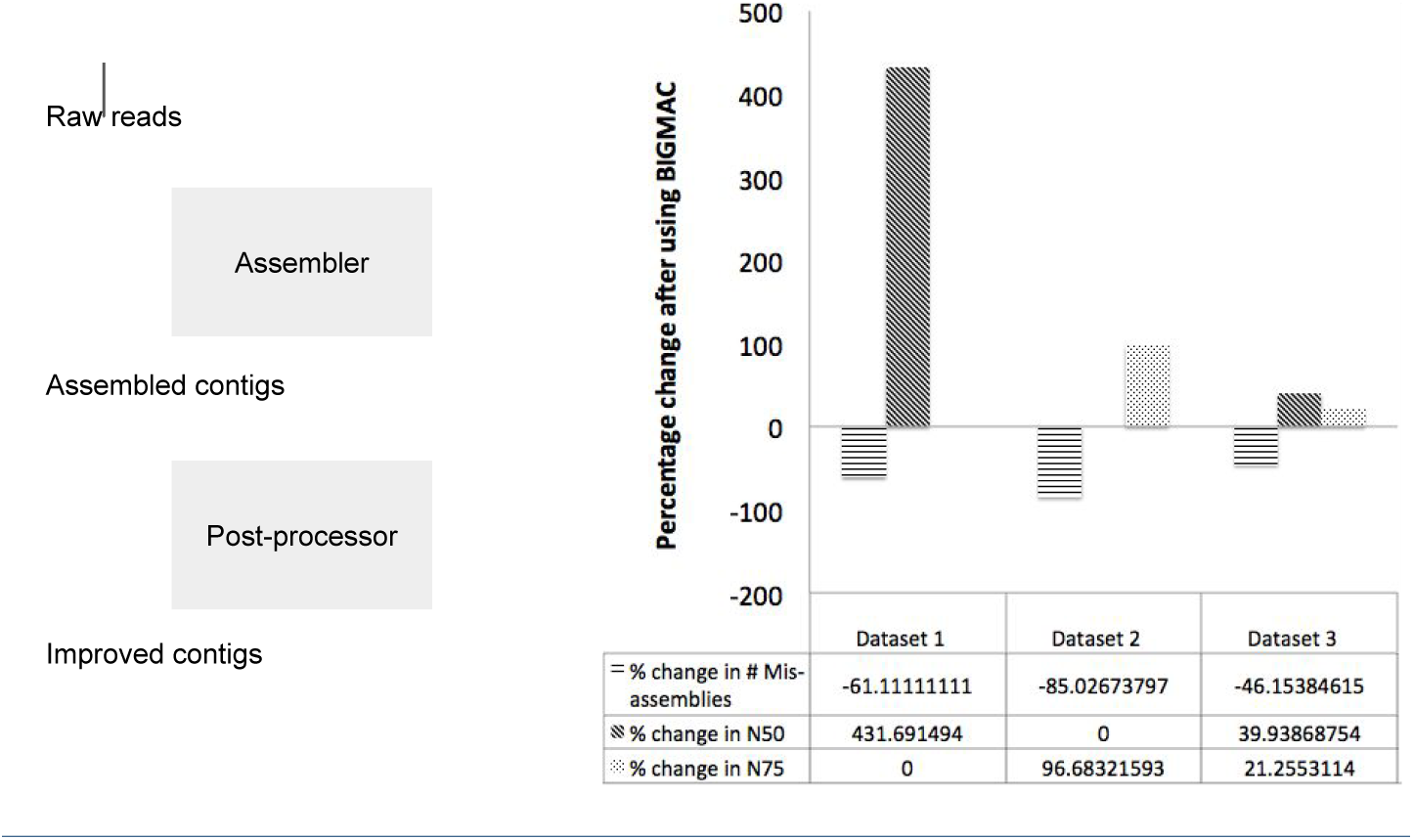
Position of post-processor in an assembly pipeline (left). Improvement in assembly quality using post-processor BIGMAC on three different datasets (right). BIGMAC shows the effectiveness of the post-processing approach for long read metagenomic assembly.

BIGMAC is a post-processor to improve metagenomic assemblies with long read only data. It first breaks contigs at potentially mis-assembled locations and subsequently scaffolds contigs. In our experiments, BIGMAC demonstrates promising results on several mock communities using data from the Pacific Biosciences long read sequencer. Inputs to BIGMAC include assembled contigs from HGAP [5] and the original raw reads with adapters removed. After assembly and post-processing, we use QUAST [6] to evaluate and compare the assembly quality of contigs before and after using BIGMAC. As shown in Fig 1, BIGMAC improves the quality of the contigs by reducing the number of mis-assemblies while maintaining/increasing N50 and N75. This demonstrates the effectiveness of the post-processing approach for metagenomic assembly with long reads.

## 2 A top-down design of BIGMAC

We use a hypothetical yet representative set of input data to illustrate the design of BIGMAC in a top-down manner. Let *g*_1_, *g*_2_ be two genomes of abundances λ_1_, λ_2_ respectively. Assume that they share a long repeat in the middle, that is, *g*_1_ = *x*_1_*ry*_1_, *g*_2_ = *x*_2_*ry*_2_. Unfortunately, an upstream assembler mis-assembles the reads and produces two contigs *c*_1_, *c*_2_ with incorrect joins at the repeat. That is, *c*_1_ = *x*_1_*ry*_2_, *c*_2_ = *x*_2_*ry*_1_. As an assembly post-processor, BIGMAC takes the mis-assembled contigs *c*_1_, *c*_2_ and original reads as input. Its goal is to reproduce *g*_1_, *g*_2_ To achieve this, we immediately recognize that there should be components for fixing mis-assemblies and scaffolding contigs. In BIGMAC, they are respectively Breaker and Merger. An illustration is given in Fig 2.

**Figure 2:**
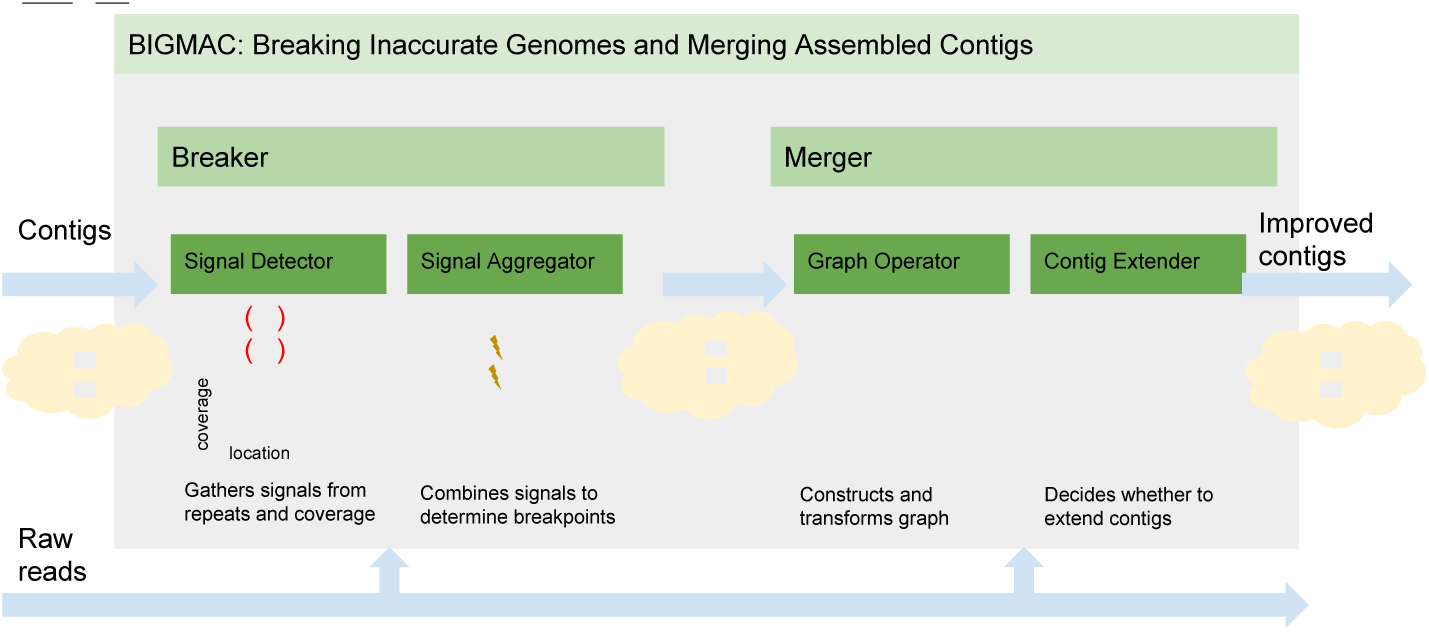
Top-down design of BIGMAC with an example of how BIGMAC improves a pair of mis-assembled contigs.

Breaker is further divided into two subcomponents: Signal Detector and Signal Aggregator. Signal Detector collects signals that indicates mis-assemblies and Signal Aggregator subsequently makes decisions on breaking contigs based on the collected signals. In our example, the coverage fluctuation between λ_1_, λ_2_ along the contigs *c*_1_, *c*_2_ and the presence of long repeat *r* are useful signals that Signal Detector collects. After aggregating these signals, Signal Aggregator decides on breaking both the contigs *c*_1_ and *c*_2_ at the starting points of the repeat *r*. Therefore, the output contigs of Breaker are *x*_1_, *x*_2_, *ry*_1_, *ry*_2_.

Merger is also divided into two subcomponents: Graph Operator and Contig Extender. With information from the original reads, Graph Operator establishes connectivity of the input contigs using string graphs. Then, based on the evidence from spanning reads and contig coverage, Contig Extender extends input contigs into longer contigs. In our example, the input contigs to Merger are *x*_1_, *x*_2_, *ry*_1_, *ry*_2_. Graph Operator forms a string graph with edges *x*_1_ → *ry*_1_, *x*_1_ → *ry*_2_, *x*_2_ → *ry*_1_ and *x*_2_ → *ry*_2_. Contig Extender analyzes the coverage depth of the related contigs and decides to merge contigs to form *ry*_1_, *x*_1_ and *ry*_2_, *x*_2_, thus reproducing the correct genomes.

## 3 Breaker: Breaking Inaccurate Genome

After the functional decomposition of BIGMAC in the previous section, we are ready to investigate its first component: Breaker. We note that the goal of Breaker is to fix mis-assemblies. In order to achieve that, we need to collect sensible signals related to mis-assemblies and subsequently aggregate the signals to make the contig breaking decisions.

### 3.1 Signal Detector

Signal Detector collects three important signals related to mis-assemblies.

#### Palindrome

We are interested in palindromes because of their relationship to a form of chimeric reads, the adaptor-skipped reads, which are common in today’s long read technology. Since assemblers get stuck at these chimeric reads, the palindrome pattern in reads propagates to the corresponding contigs. Thus, the pattern of palindrome is a strong signal indicating mis-assemblies, particularly when the palindrome is long. A string tuple (*a*, *b*) is called a wrapping pair if the reverse complement of *a* is a prefix of *b* or the reverse complement of *b* is a suffix of *a*. *x* is called a palindrome if it is the concatenation of a wrapping pair (*a*, *b*), that is *x* = *ab*. The wrapping length of *x* is max_*x*=*ab*,(_a_,*b*) is a wrapping pair_ min(*a.length*,*b.length*). For example, *x* = *ACGGCCG* is a palindrome of wrapping length 3; (*a*, *b*) = (*ACGG*, *CCG*) is a wrapping pair because the reverse complement of *b* is *CGG*, which is a suffix of *a*. Since the minimum length of *a* and *b* is min(4, 3) = 3 and the wrapping length of *x* cannot exceed 3, the wrapping length for *x* is 3.

Signal Detector collects information about palindromes by aligning each contig against itself. It then identifies palindromes with long wrapping length because that indicates mis-assemblies. The corresponding palindromes’ information is then put into *S*_*palindrome*_. To improve sensitivity, Signal Detector allows approximate match when searching for palindromes.

#### Repeat

Since long repeats confuse assemblers, their endpoints are possible candidates for mis-assemblies. Let *s*_1_ = *axb*,*s*_2_ = *cxd*, a repeat between *s*_1_, *s*_2_ is specified by the endpoints of *x* in *s*_1_, *s*_2_. For example, *s*_1_ = *CAAAAT*, *s*_2_ = *GAAAAG*, the endpoints of the repeat *AAAA* are the position specified by ! in *C*!*AAAA*!*T*, *G*!*AAAA*!*G*. Signal Detector aligns contigs against themselves to find the repeats. It then marks down the positions of the endpoints and puts them in a set called *S*_*repeat*_. To improve sensitivity, Signal Detector allows approximate matches when searching for repeats. Moreover, it only considers repeats that are maximal and are of significant length.

#### Coverage

Significant coverage variation along contigs can correspond to false joins of sequences from different genomes with different abundances. Coverage profile is the coverage depth along the contigs. For example, the coverage profile along a string *s* = *ACGT* is (1,2,2,1) if the reads are *AC*, *CG*, *GT*. Signal detector aligns original reads to the contigs to find the coverage profile, which is called *S*_*coverage*_.

### 3.2 Signal Aggregator

After Signal Detector collects signals regarding palindromes *S*_*palindrome*_, repeats *S*_*repeat*_ and coverage profile *S*_*coverage*_, Signal Aggregator uses them to determine the breakpoints on the input contigs C. The algorithm is summarized in Alg 1.

#### Algorithm 1 Signal Aggregator

1: **Input:** Input contigs *C* and signals from Signal Detector *S*_*palindrome*_, *S*_*repeat*_ and *S*_*coverage*_
2: **Output:** Contigs *C*″ with less mis-assemblies
3: **procedure** SignalAggregation(*S*_*palindrome*_, *S*_*repeat*_, *S*_*coverage*_, *C*)
4: *C*′ = ChimericContigFixing(*S*_*palindrome*_, *C*). ▹ Fix chimeric contigs
5: *S*_*filter*_ = LocatePotentialMisassemblies(*S*_*repeat*_, *C*′). ▹ Locate mis-assemblies caused by repeats
6: *C*″ = ConfirmBreakPoints(*S*_*filter*_, *S*_*coverage*_, *C*′). ▹ Confirm mis-assemblies using coverage
7: **return** *C*″

#### 3.2.1 ChimericContigFixing

The goal of ChimericContigFixing is to fix the contigs formed from chimeric reads. We simply break the palindromes in *S*_*palindrome*_ at locations corresponding to their wrapping lengths. After removing redundant contigs, ChimericContigFixing returns the broken palindromes with the unchanged contigs.

#### 3.2.2 LocatePotentialMisassemblies

The goal of the subroutine LocatePotentialMisassemblies is to locate potential mis-assemblies caused by repeats. We study the design of this subroutine in this section.

**Motivating question and example:** We can declare all the endpoints of approximate repeats in *S*_*repeat*_ to be potential mis-assemblies. While this is a sensible baseline algorithm, it is not immediately clear whether it is sufficient or necessary. It is thus natural to consider the following question.

Given a set of contigs, how can we find the smallest set of locations on contigs to break so that the broken contigs are consistent with any reasonable ground truth? To illustrate our ideas, we consider an example with contigs *x*_1_ = *abcde*, *x*_2_ = *fbcg*, *x*_3_ = *hcdi* with {*a*, *b*, *c*, *d*, *e*, *f*, *g*, *h*, *i*} being strings of equal length *L*.

The baseline algorithm of breaking contigs at the starting points of all the long(≥ 2*L*) repeats breaks the contigs 4 times(i.e. *a*|*b*|*cde*,*f*|*bcg*,*h*|*cdi*). However, interestingly, we will show that one only need to break the contigs 3 times to preserve consistency (i.e. *x*_1_ = *ab*|*cde*,*x*_2_ = *fb*|*cg*,*x*_3_ = *h*|*cdi*) and that is optimal.

**Modelling and problem formulation:** We will formalize the notions of feasible break points, feasible ground truth, consistency between sets of contigs, sufficiency of break points to achieve consistency and the optimiality criterion.

We use a graph theoretic framework. Specifically, we study a directed graph *G* = (*V*, *E*) with m sources *S* and *m* sinks *T* where ∀*υ* ∉ *S* ∪ *T*, *indeg*(*υ*) = *outdeg*(*υ*) and parallel edges between two vertices are allowed. This is used to model a fully contracted De Bruijn graph formed by successive K-mers of the contigs. Vertices *V* are substrings of the contigs and edges E correspond to potentially mis-assembled locations on contigs. In our example, the set of vertices is *V* = {*a*, *b*, *c*, *d*, *e*, *f*, *g*, *h*, *i*} and the set of edges is *E* = {*ab*, *fb*, *bc*_1_, *bc*_2_, *hc*, *cd*_1_, *cd*_2_, *cg*, *de*, *di*}. We use subscripts to differentiate parallel edges joining the same vertices. The graph corresponding to our running example is shown in Fig 3.

**Figure 3:**
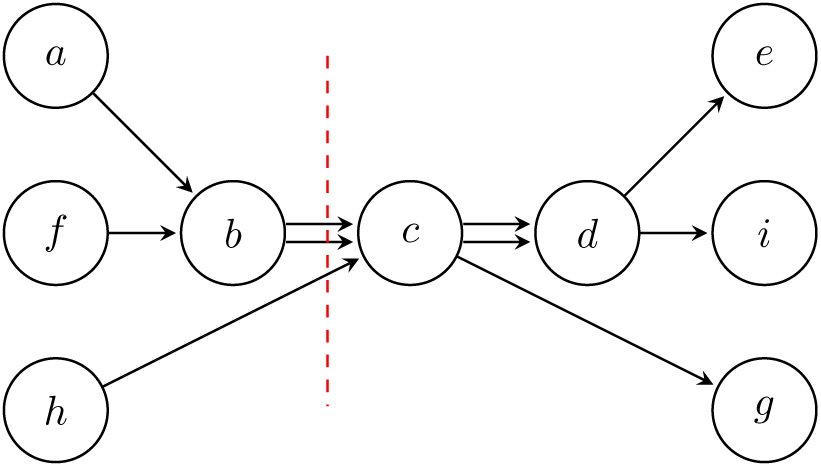
The graph corresponding to our example contig set *x*_1_ = *abcde*, *x*_2_ = *fbcg*, *x*_3_ = *hcdi* is shown. We note the optimal set of break points by the red dotted line.

Given such a graph *G*, we note that *E* is the set of all <u>feasible break points</u> because each edge in the graph corresponds to a potentially mis-assembled location on contigs. A <u>feasible ground truth</u> corresponds to a set of m edge-disjoint source-to-sink trails that partitions the edge set *E*. For simplicity, we represent a trail as a sequence of the vertices in *G*, where the edges linking the vertices are ignored.

For example, {*abcde*, *fbcdi*, *hcg*} is a feasible ground truth while {*abcg*, *fgde*, *hcdi*} is another feasible ground truth. The set of all feasible ground truths is *GT*.

We recall that our high level goal is to choose a set of feasible break points *R* ⊆ *E* so that the broken contigs are consistent with any feasible ground truth. So, we need to define the notion of broken contigs and consistency between two sets of contigs under the current framework. Let *gt* ∈ *GT*, we denote *gt*\*R* be a set of trails after the removal of the edge set *R*. In particular, for the original contig set *C* ∈ *GT*, *C*\*R* is the set of broken contigs for the set of feasible break points *R*. For example, if *R* = {*bc*_1_,*bc*_2_, *hc*} and *C* = {*abcde*, *fbcdi*, *hcg*}, *C*\*R* = {*ab*,*cde*, *fb*,*cdi*,*hcg*}. To capture consistency between two sets of contigs, we use the following definition. Given two sets of trails *T*_1_, *T*_2_, we say that *T*_1_ is <u>consistent</u> with *T*_2_ if ∀*x* ∈ *T*_1_, ∃*y* ∈ *T*_2_ *s.t*. *x* ⊆ *y* and ∀*x*′ ∈ *T*_2_, ∃*y*′ ∈ *T*_1_ *s*.*t*. *x*′ ⊆ *y*′. We call *R* a <u>sufficient breaking set</u> with respect to (*C*, *GT*) if ∀*gt* ∈ *GT*, *C*\*R* is consistent with *gt*\*R*. In other words, R is a set of feasible break points that allows the broken contigs to be consistent with any feasible ground truth. Although this notion of sufficient breaking set is a natural model of the problem, it depends on the original contig set *C*, which is indeed not necessary. Specifically, we show that we have an equivalent definition of sufficient breaking set without referring back to the original contig set. Let us call R a sufficient breaking set with respect to *G*, or simply a sufficient breaking set, if ∀*gt*_1_, *gt*_2_ ∈ *GT*, *gt*_1_\*R* is consistent with *gt*_2_\*R*.

##### Proposition 3.1

*R* is a sufficient breaking set with respect to (*C*, *GT*) if and only if *R* a sufficient breaking set with respect to *G*.

*Proof* The backward direction is immediate because *C* ∈ *GT*. We will show the forward direction as follows. Let *g*_1_, *g*_2_ ∈ *GT* and we want to show that *g*_1_\*R* is consistent with *g*_2_\*R*. Since *R* is a sufficient breaking set with respect to (*C*, *GT*), *g*_1_\*R* is consistent with *C*\*R*. Therefore, *x* ∈ *g*_1_\*R*∃*y* ∈ *C*\*R s*.*t*. *x* ⊆ *y*. But since *g*_2_\*R* is consistent with *C*\*R*, we have ∃*z* ∈ *g*_2_\*R s*.*t*. *y* ⊆ *z*. By transitivity, we have *x* ⊆ *y* ⊆ *z* ∈ *g*_2_\*R*. By symmetry, we can also show that ∀*x*′ ∈ *g*_2_\*R*∃*y*′ ∈ *g*_1_\*R s*.*t*. *x*′ ⊆ *y*′. Thus, *g*_1_\*R* is consistent with *g*_2_\*R*.

Now, we state our <u>optimization criterion</u>. The goal here is to minimize the cardinality of the sufficient breaking set, formally as Eq 1.

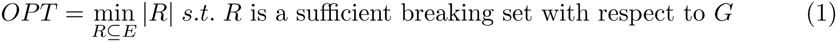

We will show that if we solve Eq 1 for our running example, the answer is 3. This thus shows that the baseline algorithm of breaking contigs at all starting points (in our example, there are 4 of them) of all long repeats is not optimal.

##### Proposition 3.2

For our running example, *OPT* = 3.

*Proof* First, by inspecting the 6 feasible ground truths in *GT*, we note *R* = {*bc*_1_, *bc*_2_, *hc*} is a sufficient breaking set with respect to *G*. Second, we note that the edge set need to disconnect sources and sinks, otherwise, we can find *g*_1_, *g*_2_ ∈ *GT* such that *g*_1_\*R*, *g*_2_\*R* are inconsistent. This means |*R*| need to be ≥ mincut of the graph, which is 3.

**Algorithm development and performance guarantee:** Next we describe an algorithm that finds a sufficient breaking set with respect to *G*. Let us denote a boolean variable *b*_*e*_ on each edge *e* ∈ *E*, with 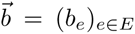. For *υ* ∈ *V*, *InEdges*(*υ*), *OutEdges*(*υ*) are the sets of incoming edges and outgoing edges at *υ* respectively. *Prev*(*υ*), *Succ*(*υ*) are the sets of predecessor vertices and successor vertices of *υ* respectively. Our algorithm solves the following minimization problem (Eq 2) as a proxy to (Eq 1).

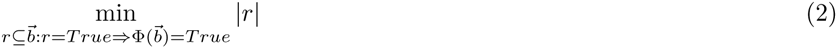

where,

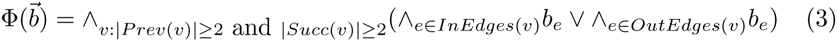

In other words, it includes either all the incoming edges or all the outgoing edges for every vertices with at least 2 successors and at least 2 predecessors to *R*. We then seek *R* with minimum cardinality among the choices available. If *G* can be decomposed into connected components, we can optimize 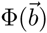 independently on each connected component. In our implementation, if the size of the connected component is not too large, we optimize the objective function by exhaustion. Beyond a certain threshold, we simply output a feasible solution without optimizing. Indeed, in our experiments on real data, most of the connected components are not that large and this step typically takes a few minutes. But we remark that for more complex applications, one can further optimize the algorithm. For example, one can first topologically sort the vertices and then use dynamic programming to solve Eq 2 along the sorted vertices.

We note that the algorithm described gives an optimal solution for our running example. Moreover, we show performance guarantee of the algorithm as follows.

##### Proposition 3.3

Solving Eq 2 gives a sufficient breaking set *R* if the graph *G* is fully contracted.

*Proof* Let *R* be the set of edges selected by the algorithm. For any two *gt*_1_, *gt*_2_ ∈ *GT*, we want to show that *gt*_1_\*R* and *gt*_2_\*R* are consistent. By symmetry, it suffices to prove that if *x* ∈ *gt*_1_\*R*, then ∃*y* ∈ *gt*_2_\*R s*.*t*. *x* ⊆ *y*. If |*x*| = 2, it is immediate because every edge other than those in R are covered. If |*x*| ≥ 3, we will show that it is also true using proof by contradiction. If ∀*y* ∈ *gt*_2_\*R*, *x* ⊈ *y*, we can find a sub-trail *x*″ = (*a*_1_, *a*_2_,…, *a*_*k*_, *a*_*k*+1_) of *x* such that ∃*y*′ ∈ *gt*_2_\*R s*.*t*. *x*″ = (*a*_1_,…, *a*_*k*_) ⊆ *y*′ but ∀*y* ∈ *gt*_2_\*R*, *x*′ ⊈ *y*. This implies ∃*a*^*^ ≠ *a*_*k* + 1_ *s*.*t*. (*x*″, *a*^*^) ⊆ *z* for some *z* ∈ *gt*_2_\*R*. Since the edge (*a*_*k*_, *a*_*k*+1_) ⊆ *x* ∈ *gt*_1_\*R*, we know that (*a*_*k*_, *a*_*k* + 1_) is not in *R*. Similarly, (*a*_*k*_, *a*^*^) ∉ *R* because (*a*_*k*_, *a*^*^) ⊆ *y*′ ∈ *gt*_2_\*R*. But since |*Succ*(*a*_*k*_)| ≥ 2, the fact that our algorithm does not include (*a*_*k*_, *a*^*^), (*a*_*k*_, *a*_*k* +1_) in *R* implies that |*Pred*(*a*_*k*_)| = 1, namely *Pred*(*a*_*k*_) = {*a*_*k*–1_}. Note that we are considering a fully contracted graph. So, the fact that *a*_*k*–1_ exists implies that |*Succ*(*a*_*k*–1_)| ≥ 2. But our algorithm has not removed edge (*a*_*k*–1_, *a*_*k*_). This gives |*Pred*(*a*_*k*–1_)| = 1. Inductively, we get |*Pred*(*a*_*i*_)| = 1∀2 ≤ *i* ≤ *k*. But we know that (*a*_*k*_, *a*_*k* + 1_) ⊆ *w* for some *w* ∈ *gt*_2_\*R*. Since the edges along (*a*_1_,…, *a*_*k*+1_) are not in *R*, this gives, *x*′ = (*a*_1_,…, *a*_*k*+1_) ⊆ *w* ∈ *gt*_2_\*R*. This contradicts the assumption about *x*′.

##### Proposition 3.4

If the graph *G* is fully contracted DAG without parallel edges, then solving Eq 2 returns a sufficient breaking set that is of optimal cardinality, OPT.

*Proof* It suffices to show that for any sufficient breaking set *R*, ∀*v* ∈ *V* where |*Succ*(*v*)| ≥ 2, |*Pred*(*v*)| ≥ 2, we have either *InEdges*(*v*) ⊆ *R* or *OutEdges*(*v*) ⊆ *R*. We prove it by contradiction. If not, ∃*v* ∈ *V* where |*Succ*(*v*)| ≥ 2, |*Pred*(*v*)| ≥ 2 but *InEdges*(*v*) ⊆ *R* and *OutEdges*(*v*) ⊆ *R*. Because it is a DAG, it means we can find *gt*_1_ ∈ *GT* such that ∃*x*, *y*, *x*′, *y*′ such that (*x*, *v*, *y*) ∈ *gt*_1_ and (*x*′, *v*, *y*′) ∈ *gt*_1_. Now consider *gt*_2_ to be a clone of *gt*_1_ except that it has (*x*, *v*, *y*′), (*x*′, *v*, *y*) instead of (*x*, *v*, *y*′), (*x*′, *v*, *y*). We note that *gt*_2_ ∈ *GT*. And because there are no parallel edges, (*x*, *v*, *y*) is not a subset of any of the trails in *gt*_2_. So, we find two distinct *gt*_1_, *gt*_2_ ∈ *GT* such that *gt*_1_, *gt*_2_ are not consistent. This contradicts the fact that R is a sufficient breaking set.

It is noteworthy that if we are given any set of contigs from any *gt* ∈ *GT*, we still obtain the same set of broken contigs after the operation of removal of redundant trails, Remove Redundant (i.e. we eliminate the contigs in a set *A* to form a minimal subset *B* ⊆ *A* in which ∀*x* ≠ *y* ∈ *B*, *x* ⊆ *y*). This can be formalized as follows.

##### Proposition 3.5

*If R is a sufficient breaking set, then for any gt*_1_, *gt*_2_ ∈ *GT, Remove Redundant (gt*_1_\*R*) = *RemoveRedundant* (*gt*_2_\*R*)

*Proof* Let *s*_*i*_ = *RemoveRedundant*(*gt*_*i*_\*R*) *for i* ∈ {1, 2}. By symmetry, it suffices to prove that *s*_1_ ⊆ *s*_2_ ∀*x* ∈ *s*_1_ ⊆ *gt*_1_\*R*, ∃*y* ∈ *gt*_2_\*R*, such that *x* ⊆ *y*. Note that we can also find some *x*^*^ ∈ *s*_2_ such that *y* ⊆ *x*^*^. This gives *x* ⊆ *y* ⊆ *x*^*^. However, since we have no redundant trails in *s*_1_, we get *x* = *x*^*^. Thus *x* = *x*_*_ ∈ *s*_2_.

To apply BIGMAC to real data, we have to implement the described algorithm with some further engineering. This includes methods to tolerate noise, to handle double stranded nature of DNA, and to construct De Bruijn graph from the repeats. Interestd readers can refer to the Appendix for these implementation details.

#### 3.2.3 ConfirmBreakPoints

The goal of ConfirmBreakPoints is to utilize the coverage profile *S*_*coverage*_ to confirm breaking decisions at potentially mis-assembled locations specified in *S*_*filter*_. For the sake of simplicity, we now consider a string *s* of length *L*, and a set of positions *Y* = {*y*_*i*_}_1≤*i*≤k_ of *s* which are potential mis-assemblies. Furthermore, let us assume that all mis-assemblies are caused by mixing genomes of different abundances. Using *Y*, we can partition s into segments {*s*_*i*_}_0≤*i*≤k_ of lengths respectively as {*ℓ*}_1≤*i*≤k_ of *s*. We let *x*_*i*_ be the number of reads that start in segment *s*_*i*_, which can be estimated from *S*_*coverage*_. The question is how to classify the points in *Y* as true mis-assemblies or as fake mis-assemblies.

One can use heuristics, like comparing coverage depth difference before and after each *y*_*i*_. However, this is not ideal. For example, if we have coverage depth before and after *y*_1_ differing by 1X, we would expect it to be a much stronger signal for true mis-assembly if the lengths *ℓ*_0_, *ℓ*_1_ are of order of 100K rather than of 100 and this cannot be seen by considering coverage depth difference alone. For the case of just two segments of equal length and if we assume the *x*_*i*_’s are independent Poisson random variables, there is a popular statistical test, C-Test[7], that can make the decision. Formally, if *x*_1_ ~ *Poisson*(*m*_1_) and *x*_2_ ~ *Poisson*(*m*_2_), then C-Test is a test to decide between the hypothesis of *H*_0_: *m*_1_ = *m*_2_ against *H*_1_: *m*_1_ = *m*_2_. C-Test first considers *x*_1_ + *x*_2_ to compute the decision boundary and it then decides whether to reject *H*_0_ based on *x*_1_/*x*_2_ and the previously derived decision boundary. The intuition is that *x*_1_ + *x*_2_ corresponds to the amount of data, which determines the confidence of a statistical test. Thus, if *x*_1_ + *x*_2_ is large, a small perturbation of *x*_1_/*x*_2_ from 1 can still be very significant and can correspond to a confident rejection of *H*_0_.

However, we still need to think carefully about how to apply C-Test to our problem. One method is to directly apply k independent C-Test on each of the junctions *y*_*i*_. However, that method cannot take advantage of the information from neighboring segments to boost the statistical power at a junction. Therefore, we develop the algorithm IterativeCTest. IterativeCTest performs traditional C-Test but in multiple iterations. Initially, it directly applies *k* independent C-Test on each of the junctions *y*_*i*_. Some of the segments are merged and we aggregate the counts from the merged segments to continue to the next C-Test at the remaining unmerged junctions in *Y*. This process is repeated until the algorithm converges. Finally, we use the breaking decisions from IterativeCTest to break the incoming contigs and return the fixed contigs.

## 4 Merger: Merging Assembled Contigs

After fixing mis-assemblies, we are ready to study another pillar of BIGMAC: Merger. Merger establishes connectivity among contigs and subsequently makes decisions to extend contigs.

### 4.1 Graph Operator

The goal of Graph Operator is to identify candidates for contig extension. It forms and transforms a string graph using the overlap information among original reads and contigs. It subsequently analyzes the graph to find candidates for contig extension. The overall algorithm is summarized in Alg 2.

**Mapping:** We obtain mapping among contigs and reads. This provides the fundamental building block to construct the connectivity relationship among contigs and reads.

**FormGraph:** The goal of FormGraph is to establish connectivity among contigs. We use contig-read string graph as our primary data structure. Contig-read string graph is a string graph[8] with both the contigs and the reads associated with their endpoints as nodes. The size of the graph is thus manageable because most reads are not included in the graph. Then, we construct an overlay graph (which we call the contig graph) such that the nodes are the contigs with weights on nodes being the coverage depth of contigs. In the contig graph, there is an edge between two nodes if there is a sequence of reads between the associated contigs. With these data structures, we can switch between macroscopic and microscopic representation of the contig connectivity easily.

#### Algorithm 2 Graph Operator

1: **Input:** Contigs *C* and original reads *R*
2: **Output:** String graph *G* with information about candidates for contig extension
3: **procedure** GraphOperator(*R*, *C*)
4: M = Mapping(R, S). ▹ Obtain mapping among contigs and reads
5: G = FormGraph(M). ▹ Form string graph to represent connectivity
6: G.GraphSurgery(M). ▹ Simplify graph
7: G.FindExtensionCandidates(). ▹ Identify candidates for contig extension
8: **return** G

**GraphSurgery:** GraphSurgery simplifies the contig graph. This includes removal of transitive edge and edge contraction.

For nodes *u*, *v*, *w*, if we have edges *u* → *v*, *u* → *w* and *w* → *v*, then we call *u* → *v* a transitive edge. We perform transitive reduction[8] on the contig graph to remove transitive edges. Removing these transitive edges prevents us from finding false repeats in the next stage. To improve robustness, there is a pre-processing step before transitive reduction. If the contig *w* is too short and there are no reads that form head-to-tail overlap between *w*, *u* or *w*, *v*, then we can have a missing edge for transitive reduction to operate properly. Thus, we add the missing edge (either from *u* to *w* or *w* to *v*) back when we find contigs and related reads that suggest the missing edge might be there.

An edge *u* → *v* is contractable when the outgoing degree of *u* and the incoming degree of v are both 1. We contract edges to fill gaps. Our experience with FinisherSC is that data are dropped in the assembler and so reconsidering them as a post-processing step can potentially fill some gaps. However, there is a caveat. In establishing connectivity in contig-read string graph, we only use reads close to each contig’s endpoints (as a way to save computation resources), we may miss some legitimate connections. Therefore, very long repeats prevent the detection of linkage of contigs, thereby allow contigs to be erroneously merged. To address that, if there exists contig *w* that is connected (by some reads) to a region close to the right end of *u*/left end of *v*, then we avoid contraction of *u* → *v*.

**FindExtensionCandidates:** FindExtensionCandidates identifies candidates for contig extension by identifying local patterns in the contig graph. One form of extension candidates is a pair of contigs that are connected without competing partners. This corresponds to the contigs joined by a contractable edge. Another form of extension is a set of contigs that are connected with competing partners. This corresponds to the contigs linked together in the presence of repeats. In the contig graph, the repeat interior can either be represented as a separate node or not. If the repeat interior is represented as a separate node, the local subgraph is a star graph with the repeat node at the center. Otherwise, the local subgraph is a bipartite graph consisting of competing contigs. After identifying the contigs associated with a specific repeat, we can then merge contigs in the next stage.

### 4.2 Contig Extender

After the operations by Graph Operator, we have extracted the potential contig extension candidates from the contig graph. It remains to decide whether and how to merge the corresponding contigs. In a high level, Contig Extender aims at solving the Contig Merging Decision Problem.

**Contig Merging Decision Problem** *Given a set of incoming contigs I and a set of outgoing contigs O whose coverage depth and read connectivity information is known. Decide how to form an appropriate bipartite matching between I and O*.

While we do not intend to rigorously define the statement of Contig Merging Decision Problem now, it is clear that appropriate matching corresponds to one that achieves high accuracy. Contig Extender carefully analyzes the read connectivity and contig coverage to make decisions on merging contigs. In the coming discussion, we first study an effective heuristic that captures the essence of the problem. After that, we will study how to rigorously define the Contig Merging Decision Problem in a mathematical form and suggest an EM-algorithm in solving that.

#### 4.2.1 Heuristic in solving the Contig Merging Decision Problem

When the cardinality of the set of incoming contigs I and the set of outgoing contigs *O* are both 1, a natural decision is to merge them. Thus, the focus here is to study the case when |*I*| > 1 or |*O*| > 1. We select the reads that uniquely span one contig a in the incoming set and one contig *b* in the outgoing set. If there are multiple such reads, then we decide that *a*, *b* should be joined together provided that there does not exist any paths of reads that lead *a* to other contigs in the outgoing set and similarly for *b*. Moreover, we perform similar tests using contig coverage. If the coverage depth between two contigs is very close, they will be declared to be a potential merge pair. Then, we test whether there are any close runner-ups. If not, they should be merged. To combine the decisions made using spanning reads and coverage depth, we have a subroutine that discards all conflicting merges. We find that this heuristic for decision making works quite well in our datasets. However, in the coming discussion, we will study how to make the extension decisions in a more principled and unified manner.

#### 4.2.2 General framework in solving the Contig Merging Decision Problem

The challenge for the Contig Merging Decision Problem is the tradeoff for many physical quantities (e.g. abundance, edit distance of reads, noise level, number of spanning reads, etc). We address this by defining a confidence score based on parameter estimation. For simplicity of discussion, we assume that *k* is the cardinality of both the set of incoming contigs and the set of outgoing contigs. The goal is to find the best perfect matching with respect to a confidence score.

Let *M* be a perfect matching of contigs among incoming and outgoing groups *I* and *O*. Under *M*, there are *k* merged contigs. Let the observation be the set of related reads *X* = {*R*_*i*_ | *1* ≤ *i* ≤ *n*}. We define the parameters *θ* = {(*λ*_*j*_, *I*_*j*_)_1≤*j*≤*k*_}, where *λ*_*j*_ is the normalized abundance of contig *j* and *I*_*j*_ is genomic content of the contig *j*. Note that Σ_1≥*j*≥*k*_ *λ*_*j*_ = 1. So, the parameter estimation problem can be cast as *s*_*M*_ = max_*θ*_ *P*_*θ*_(*X* | *M*), where *s*_*M*_ is the confidence score of the matching *M*. Finally, the best perfect matching can be found by comparing the corresponding confidence score.

Note that we have hidden variables *Z* = (*Z*_*i*_)_1≤*i*≤*n*_ which indicate the contigs that reads *X* are extracted from (i.e. *Z*_*i*_ ∈ {1, 2,…,*k*}). If we assume the length of the contig *j* to be *ℓ*_*j*_ and *q* to be the indel noise level (i.e. probability of 1 – 2*q* to be remained unaltered at each location), then we can use an EM-algorithm to obtain an estimate of *θ*. Specifically, the expected value of the log likelihood function *E*_*q*(*Z*|*X*, *θ*^(*t*)^)_ [log *P*_*θ*^(*t*)^_ (*X*, *Z*, *θ*^(*t* + 1)^] is

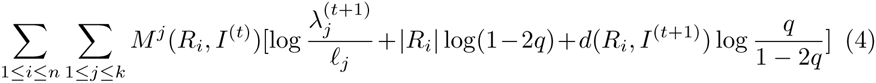

where 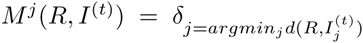 is the assignment of reads to the most similar contig (with tie breaking using *λ*^(*t*)^), *d*(*A*, *B*) is the best local alignment score, 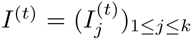 are the genomic contents of the contigs at iteration *t* and *λ*^(*t*)^ = (*λ*_*j*_)_1≤*j*≥*k*_ are the estimated abundances at iteration *t*. By maximizing the log likelihood function with respect to *θ*^(*t*+1)^, we have the update formulas as

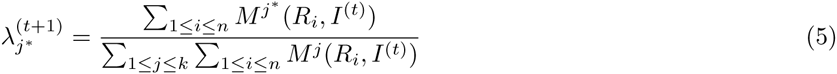

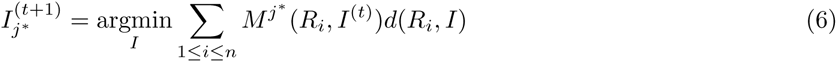

Note that the update of *I*_*j*\*_ requires multiple sequence alignment. In general, the problem is NP-hard[9]. However, for noisy reads extracted from several contigs, the problem is not as difficult. For example, in the case of pure substitution noise, an efficient optimal solution is a column-wise majority vote. Despite the elegance and feasibility of this approach, it is still computationally more intense than the heuristic. Therefore, in our implementation of BIGMAC, the heuristic is the default method used in Contig Extender. But we also provide an experimental version of the EM-algorithm which can be used when users specify `--option emalgo=True` in their commands.

## 5 Experiments

### 5.1 End-to-end experiments

#### 5.1.1 Synthetic data

We verify that BIGMAC correctly improves the incoming contigs using the following synthetic data. We generate two synthetic species of different abundances 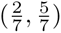. They are composed of random nucleotide sequences of length 5M each and share a common segment of length 12K. We uniformly sample indel noise corrupted reads of length 6K from the genomes, with coverage depth of 20X and 50X respectively. We also artificially construct mis-assembled contigs by switching the genome segments at the 12K repeat.

The contigs and the reads are passed through BIGMAC. BIGMAC breaks the contigs at the mis-assembled point and joins them back correctly. One can see the sample run on [10]. This is also the example that we discuss in the top-down design of BIGMAC.

#### 5.1.2 Real data

We test the performance of BIGMAC in improving metagenomic assembly on PacBio only data. We use different datasets of different characteristics. Dataset 1 consists of a mock community of 5 species [11], with genomes of high similarity. Dataset 2 consists of a mock community of 23 species [12], with genomes of diverse abundances. We also remark that we have tested BIGMAC on a third PacBio only dataset (Dataset 3): a real experiment involving Desulfuromonas biwabikus, D. soudanensis and some other unknown genomes. We note that we know the complete ground truth for the metagenomes in both Dataset 1 and 2 but only know part of the ground truth for Dataset 3. We take the output of HGAP and use the raw reads to improve them using BIGMAC. We observe that in these datasets, BIGMAC reduces the number of mis-assemblies while maintaining/increasing N50 and N75. The results of BIGMAC is summarized in Table 1, where the quantity of mis-assemblies is obtained from the QUAST reports. By default, QUAST’s statistics are based on contigs of size >= 500 bp. Interested readers can refer to the Appendix for more details of the assessment. The data is provided in our online distribution and users can reproduce the results by issuing a single command `python reproduce.py`

**Table 1:** Performance evaluation of BIGMAC on several mock communities is shown. BIGMAC consistently improves assembly quality by simultaneously increasing contig contiguity and decreasing the number of mis-assemblies.

| Dataset | Method | NContig | # Mis-assembly | N50 | N75 |
| --- | --- | --- | --- | --- | --- |
| 1 | HGAP | 130 | 18 | 818655 | 274801 |
| 1 | BIGMAC | 129 | 7 | 4352719 | 274801 |
| 2 | HGAP | 477 | 187 | 397611 | 38471 |
| 2 | BIGMAC | 344 | 28 | 397611 | 75666 |
| 3 | HGAP | 185 | 26 | 257044 | 82370 |
| 3 | BIGMAC | 140 | 14 | 359704 | 99878 |

### 5.2 Comparison with other post-processing tools

#### 5.2.1 Synthetic data

We use the synthetic data in Section 5.1.1 to evaluate and benchmark BIGMAC, FinisherSC[13], SSPACE_LongRead[14] and PBJelly[15]. BIGMAC is the only tool among the tested tools that fix the mis-assembled contigs and merge them back correctly. Other tested tools output the same mis-assembled contigs.

#### 5.2.2 Real data

We perform end-to-end testing to compare performance of different tools. The comparison is shown in Table 2. BIGMAC shows the largest N75/# Mis-assemblies on all three datasets and largest N50/# Mis-assemblies on two out of three datasets. Indeed, in the only dataset that BIGMAC does not have the largest N50/#Mis-assemblies, the number of contigs(i.e. L50) beyond N50 is 7. Due to the small number of contigs, this particular measurement on that dataset may not be significant. We also remark that BIGMAC is the only tool that improves contiguity (N50 and N75) and the number of mis-assemblies.

**Table 2.** Comparison of performance of BIGMAC with other post-processing tools for long read assemblies is shown. BIGMAC shows the largest N75/# Mis on all three datasets and largest N50/# Mis on two out of three datasets. We also remark that BIGMAC is the only tool that can improve N50 and N75 while reducing the number of mis-assemblies. Note that # Mis is an abbreviation for the number of mis-assemblies.

| Data | Method | # Mis | N50 | N75 | N50/# Mis | N75/# Mis |
| --- | --- | --- | --- | --- | --- | --- |
| 1 | HGAP | 18 | 818655 | 274801 | 45481 | 15267 |
|  | BIGMAC | 7 | 4352719 | 274801 | **621817** | **39257** |
|  | FinisherSC | 32 | 2531294 | 415024 | 79103 | 12970 |
|  | PBJelly | 19 | 4642330 | 418480 | 244333 | 22025 |
|  | SSPACE.LR | 32 | 4657611 | 493683 | 145550 | 15428 |
| 2 | HGAP | 187 | 397611 | 38471 | 2126 | 206 |
|  | BIGMAC | 28 | 397611 | 75666 | **14200** | **2702** |
|  | FinisherSC | 192 | 654163 | 43018 | 3407 | 224 |
|  | PBJelly | 271 | 1585584 | 61775 | 5851 | 228 |
|  | SSPACE.LR | 255 | 1568442 | 95133 | 6151 | 373 |
| 3 | HGAP | 26 | 257044 | 82370 | 9886 | 3168 |
|  | BIGMAC | 14 | 359704 | 99878 | 25693 | **7134** |
|  | FinisherSC | 25 | 996532 | 97964 | 39861 | 3919 |
|  | PBJelly | 27 | 1103847 | 128718 | **40883** | 4767 |
|  | SSPACE.LR | 43 | 1266912 | 290104 | 29463 | 6747 |

### 5.3 Simulation and testing on independent components

We perform micro-benchmarking on individual components of BIGMAC. The settings and results are summarized as follows.

Analysis of LocatePotentialMisassemblies: We test our break point finding algorithm used in LocatePotentialMisassemblies of Breaker on the synthetic data of *x*_1_ = *abcde*,*x*_2_ = *fbcg*, *x*_3_ = *hcdi* discussed in the previous section. The algorithm succeeds in identifying the minimum number of break points as 3. Also, it succeeds in identifying the minimum number of break points as 2 in the presence of double stranded DNA, in the test case of *x*_1_ = *abcd*, *x*_*2*_ = *ec*′*b*′*f*, where *b*′, *c*′ are the reverse complement of *b*, *c* respectively.

**Analysis of ConfirmBreakPoints**: We test IterativeCTest used in ConfirmBreakPoints of Breaker on synthetic data. We simulate mis-assemblies and variation on abundances by generating a sequence of Poisson random variables and compare the performance of the algorithms on the simulated data as follows. We generate a sequence of 100 numbers by 100 independent Poisson random variables. The Poisson random variables have parameters of either 20 or 50. To select the parameters, we simulate 100 steps of a two-states Markov chain with transition probability of 0.1. We then evaluate the performance of C-Test and IterativeCTest on finding the true transition points, which correspond to the junctions of mis-assemblies. Taking average from 100 rounds of simulation, the recall of both C-Test and IterativeCTest are 0.93, meaning that they both can identify almost all transition points. C-Test has precision of 0.75 while the precision of IterativeCTest is of 0.87, meaning that IterativeCTest can avoid more fake mis-assemblies.

Analysis of Merger: We compare Merger with other tools on synthetic data as follows. We use a synthetic contig set {*x*_1_, *x*_2_, *r*, *y*_1_, *y*_2_} where the ground truth genomes are (*x*_1_, *r*, *y*_1_), (*x*_2_, *r*, *y*_2_). The coverage depth of (*x*_1_, *y*_1_) and (*x*_2_, *y*_2_) are 20X and 50X respectively. We pass the reads together with the contig set to FinisherSC, PBJelly, SSPACE_LongRead to perform scaffolding. We note that BIGMAC is the only tool the can scaffold the contigs correctly into 2 contigs by using the abundance information among the tested tools. Other tools either do not change the input or simply merge r with some of {*x*_1_, *x*_2_, *y*_1_, *y*_2_}, resulting in 4 contigs.

#### 5.3.1 Runtime of BIGMAC

The runtime of BIGMAC is summarized in Table 3. We use 20 threads to run the tool on a server computer. The server computer is equipped with 64 AMD Opteron(tm) Processor 6380(8 cores) with frequency 2500 MHz and 362GB RAM. We note that the majority of the time is spent on alignment of contigs and reads by MUMmer.

**Table 3.** Runtime of BIGMAC and the corresponding file size

| Dataset | Component | Contig file size (MB) | Read file size (GB) | Running time (sec) |
| --- | --- | --- | --- | --- |
| Synthetic | Breaker | 9.6 | 0.335 | 164 |
| Synthetic | Merger | 9.6 | 0.335 | 123 |
| 1 | Breaker | 30 | 5.7 | 6646 |
| 1 | Merger | 29 | 5.7 | 6998 |
| 2 | Breaker | 32 | 5.8 | 4865 |
| 2 | Merger | 29 | 5.8 | 5087 |
| 3 | Breaker | 17 | 7.6 | 7099 |
| 3 | Merger | 14 | 7.6 | 6887 |

### 5.4 Discussion

There are two main implications from the experiments performed. First, we show that post-processing assemblies is a feasible alternative in improving assembly quality to building another assembler from scratch. This is demonstrated by BIGMAC showing simultanous improvement in terms of number of mis-assembly and contiguity. We note that this is a non-trivial feature because all other tested tools fail to achieve it. Second, BIGMAC is competitive when compared to the existing post-processing tools. In particular, it shows better N75/# Mis-assemblies than all other tested tools in all tested datasets. Moreover, BIGMAC is also the only tool that can handle abundance information, which makes it an attractive candidate for improving metagenomic assembly.

## References

1. Chen, K., Pachter, L.: Bioinformatics for whole-genome shotgun sequencing of microbial communities. PLoS Comput Biol 1(2), 106–112 (2005)

2. The Critical Assessment of Metagenome Interpretation (CAMI) competition. http://blogs.nature.com/methagora/2014/06/the–critical–assessment–of–metagenome–interpretation–cami–competition.html

3. Koren, S., Phillippy, A.M.: One chromosome, one contig: complete microbial genomes from long-read sequencing and assembly. Current opinion in microbiology 23, 110–120 (2015)

4. Lam, K.-K., LaButti, K., Khalak, A., Tse, D.: Finishersc: a repeat-aware tool for upgrading de novo assembly using long reads. Bioinformatics 31(19), 3207–3209 (2015). doi:10.1093/bioinformatics/btv280. http://bioinformatics.oxfordjournals.org/content/31/19/3207.full.pdf+html

5. Chin, C.-S., Alexander, D.H., Marks, P., Klammer, A.A., Drake, J., Heiner, C., Clum, A., Copeland, A., Huddleston, J., Eichler, E.E., et al.: Nonhybrid, finished microbial genome assemblies from long-read smrt sequencing data. Nature methods 10(6), 563–569 (2013)

6. Gurevich, A., Saveliev, V., Vyahhi, N., Tesler, G.: Quast: quality assessment tool for genome assemblies. Bioinformatics 29(8), 1072–1075 (2013)

7. Przyborowski, J., Wilenski, H.: Homogeneity of results in testing samples from poisson series: With an application to testing clover seed for dodder. Biometrika, 313–323 (1940)

8. Myers, E.W.: The fragment assembly string graph. Bioinformatics 21(suppl 2), 79–85 (2005)

9. Elias, I.: Settling the intractability of multiple alignment. Journal of Computational Biology 13(7), 1323–1339 (2006)

10. Documentation of MetaFinisherSC. https://github.com/kakitone/MetaFinisherSC

11. Hall, R.J., Chin, C.-S., Mehrotra, S., Juretic, N., Wasserscheid, J., Dewar, K.: An interactive workflow for the analysis of contigs from the metagenomic shotgun assembly of SMRT Sequencing data. http://files.pacb.com/pdf/RHall-ASM2014InteractiveWorkflow.pdf (2014)

12. PacBio: PacBio Devnet. https://github.com/PacificBiosciences/DevNet/wiki/Human_Microbiome_Project_MockB_Shotgun

13. Lam, K.-K., LaButti, K., Khalak, A., Tse, D.: Finishersc: a repeat-aware tool for upgrading de novo assembly using long reads. Bioinformatics 31(19), 3207–3209 (2015)

14. Boetzer, M., Pirovano, W.: Sspace-longread: scaffolding bacterial draft genomes using long read sequence information. BMC bioinformatics 15(1), 1 (2014)

15. English, A.C., Richards, S., Han, Y., Wang, M., Vee, V., Qu, J., Qin, X., Muzny, D.M., Reid, J.G., Worley, K.C., et al.: Mind the gap: upgrading genomes with pacific biosciences rs long-read sequencing technology. PloS one 7(11), 47768 (2012)

16. Larkin, M.A., Blackshields, G., Brown, N., Chenna, R., McGettigan, P.A., McWilliam, H., Valentin, F., Wallace, I.M., Wilm, A., Lopez, R., et al.: Clustal w and clustal x version 2.0. Bioinformatics 23(21), 2947–2948 (2007)

